# Genome-wide assessment of runs of homozygosity in Sahiwal cattle of Pakistan

**DOI:** 10.1101/2023.11.17.567615

**Authors:** Abdul Rahman Sesay, Mohammad Saif-ur-Rehman, Faisal Ramzan, Faisal Saeed Awan

## Abstract

Runs of homozygosity (ROH) are extensive stretches of homozygous genotypes present in an individual genome inherited from each of its parents. Selection pressures have been one of the core causes of a reduction in genetic diversity in some genomic regions, leading to an accumulation of ROH in these regions. An individual’s ROH can provide information about the past of his population, including the extent of inbreeding, recent bottlenecks, and positive selection. The study aimed to evaluate the degree of autozygosity throughout the genome in Sahiwal cattle to discern and describe ROH patterns. In addition, we also estimated the rate of inbreeding based on ROH in Sahiwal cattle. A sample of 98 Sahiwal bulls from various public institutional herds and private livestock farms in the Punjab province of Pakistan was used for this study. All animals were genotyped using the 140k BovineHD SNP chip. The study identified ROH in all animals. The mean ROH per animal was 29.71, with values extending from 2.35 to 85.31. The mean length of the ROH was 5.84 Mb, and the longest segment was 51.43 Mb (1,727 SNPs) found on BTA3. Results revealed that genome-wide ROH for Sahiwal was typically composed of many short and medium segments (2–4 Mb and 4–8 Mb), accounting for approximately 54.18% of all detected ROH. The inbreeding coefficient based on ROH (F_ROH_) ranged from 0.016 to 0.057. The study revealed several genes, including STAT1, ATP1A1, OLR1, and CD4, which are known genes related to milk production. Thus, understanding ROH patterns, an inbreeding coefficient derived from ROH, and candidate genes associated with significant economic traits can help explain the mechanisms governing these traits in Sahiwal cattle.

## Introduction

Runs of homozygosity (ROH) are extensive stretches of homozygous genotypes present in an individual genome inherited from each of its parents [1–3]. Loci in the genome with a common ancestral origin, known as identical by descent (IBD), are considered autozygous when passed down from both parents to their offspring [4,5]. ROH offers valuable insights into individuals’ genetic relatedness, mitigating inbreeding and uncovering detrimental genetic variations within the genome [6,7]. The genomic characteristics of runs of homozygosity (ROH) are subject to various influences, including natural as well as artificial selection, recombination patterns, linkage disequilibrium, population structure, mutation rate, as well as levels of inbreeding [5,8–10]. [11] first identified long homozygous stretches of the human genome using microsatellite markers. High-throughput genomic analysis tools, such as next-generation sequencing and genotype-based microarrays, evolved rapidly over the last 20 years, allowing ROH to be detected accurately [7,12].

ROH can produce accurate evaluations of genomic autozygosity and allows one to estimate the genomic inbreeding coefficient (F_ROH_) [13,14]. Estimates of F_ROH_ can be made for any animal with genotypic data and provide a more accurate prediction of the genome and the actual degree of autozygosity [15,16]. The utilization of genetic information from runs of homozygosity (ROH) for the computation of the inbreeding coefficient (FROH) is deemed to be a superior method in comparison to estimations derived from pedigree data (FPED) to accurately assess autozygosity and identify both historical and more recent inbreeding impacts [5]. Heterozygosity-fitness correlations (HFCs) have been broadly employed to investigate the influence of inbreeding on the fitness of individuals [17,18]. The measurement of inbreeding variance can be achieved by utilizing genetic markers, specifically by assessing the correlation strength in heterozygosity across marker loci, which is referred to as identity disequilibrium (ID) [19]. The quantification of ID can be achieved by utilizing the g2 measure, which serves as a pivotal parameter in the theory of HFC. This measure can be employed within a broader context to assess the direct influence of inbreeding on marker heterozygosity as well as fitness [20–22]. An alternative method for estimating ID involves partitioning the marker panel into two subcategories in a random manner. The correlation in heterozygosity between these subsets is then calculated, and this process is repeated numerous times to generate a distribution of heterozygosity-heterozygosity correlation (HHC) coefficients [23]. The methodology mentioned above is characterized by its intuitive nature and is comparable to g2 in terms of its ability to identify the presence of non-zero variation in inbreeding [17,18].

The Sahiwal cattle breed is Pakistan’s most important breed of dairy cattle due to its tolerance to heat, disease resistance, and adequate performance in low-quality roughages [24,25]. The practice of pure breeding, combined with selective breeding, has been extensively employed in Pakistan for the past 30 years to enhance and protect Sahiwal cattle breeds. Significant advances have been made in improving physical attributes and capabilities. However, efficiency has not improved much over the years. The impaired situation has been attributed to several factors, including inbreeding [26,27]. Inbreeding is a phenomenon that arises when animals are bred with one another despite sharing common ancestors, and it is an inevitable occurrence within commercial breeding programs specifically designed for dairy cattle [28–30]. The ramifications of inbreeding encompass genetic drift, reduction in heterozygosity, and diminishment of genetic diversity [31]. A growing focus has recently been on understanding genetic variability within populations [32–35]. The preservation of genetic diversity in the Sahiwal cattle population poses a significant challenge [36,37]. Therefore, the study aims to evaluate the degree of autozygosity throughout the genome in Sahiwal cattle to discern and describe ROH patterns. In addition, to estimate the rate of inbreeding based on ROH in Sahiwal cattle.

## Materials and Methods

### Animal resources and SNP genotyping

The Sahiwal (Bos indicus) dairy cattle bulls maintained at various public institutional herds and private livestock farms in the Punjab province of Pakistan were selected for this study. Blood samples were taken from each of the 98 Sahiwal cattle bulls. The utilization of animals and the collection of blood samples were conducted in accordance with the protocols established by the Ethics Committee of the University of Agriculture, Faisalabad. A salting-out approach was used to recover genomic DNA from blood samples [38]. The NanoDrop ND-1000 spectrophotometer was used to measure the extracted DNA concentration (NanoDrop Technologies Wilmington, DE). Following the normal operating procedures suggested by the manufacturer, the genetic makeup of each animal was determined by genotyping them using a 140k BovineHD BeadChip (Illumina Inc. in San Diego, California, USA). The PLINK program [39] performed quality assurance checks on genotyping data. The study selected a call rate greater than 95% for study data and analysis. The study used SNPs with minor allele frequencies (MAFs) less than 0.05. The study analysis retained markers and animals without a significant deviation from Hardy-Weinberg proportions (P > 0.001, Bonferroni corrected). Only SNPs found in autosomes were included in the analysis. In addition, samples with more than 10% missing genotypes were eliminated. SNPs were filtered to omit loci assigned to unmapped contigs as well as chromosomes of the sex. Following quality control, 87 samples and 74,070 SNPs were retained in 29 autosomes.

### Detection and classification of runs of homozygosity

PLINK software [39] was used to calculate each individual’s runs of homozygosity (ROH). As recommended by Purfield et al. (2012), a minimum ROH length threshold of 1 Mb was established to ensure the exclusion of short and common ROH deriving from linkage disequilibrium (LD). The PLINK parameters and thresholds specified by Purcell et al. (2007) were used to define ROH. A sliding window of 50 single nucleotide polymorphisms (SNP) covered the entire genome. The number of Overlapping windows with many homozygous genotypes ≥ 0.05 was considered. An ROH was defined as 100 or more consecutive SNPs with homozygous genotypes. The minimum ROH length has been specified at 1MB. A maximum of 100 kb was allowed between homozygous SNPs. An SNP per 50 kb was specified as the density. An ROH could contain up to five SNPs lacking genotypes and one heterozygous genotype. According to the nomenclature proposed by [41] and [6], all identified ROHs were classified into five distinct classes: 1–2 Mb, 2–4 Mb, 4–8 Mb, 8–16 Mb, and >16 Mb. Furthermore, for every individual and each ROH length category, the mean number of ROH per individual, the average length of ROH, and the total number of ROH per breed were estimated.

### Identification of common runs of homozygosity and gene annotation

The genomic areas predominantly associated with runs of homozygosity (ROHs) were determined through the computation of the SNP proportion inside ROHs. Quantification of SNP occurrence inside the regions was achieved through a systematic tallying of its frequency in many individuals. Subsequently, we identified the uppermost 1% of single nucleotide polymorphisms (SNPs) that were frequently detected inside run of homozygosity (ROHs). SNPs that were found to be in close proximity and exceeding the specified threshold were eventually combined to form genomic regions known as ROH islands. These ROH islands are distinguished by their prevalence among the majority of individuals within the population [42]. The ARS-UCD1.2 assembly databases [43] and the Ensembl genome browser [44] were utilized for gene annotation in the ROH island. The QTL Animal database (available at https://www.animalgenome.org/cgi-bin/QTLdb/BT/index) [45] was used to determine the quantitative trait loci (QTL) reported in the literature for each specific candidate region. The biological function of each gene identified in the region of ROH island was deduced by conducting thorough and precise literature searches.

### Genomic inbreeding coefficients

The PLINK software was used to calculate the genomic inbreeding coefficients (F_ROH_). According to McQuillan et al. (2008), the following formula was used to determine each animal’s inbreeding coefficient based on ROH (F_ROH_):

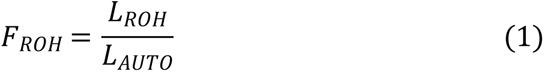

Where L_ROH_ is the entire length of ROH in the genome of individuals, while L_AUTO_ is the length of the autosomal genome that SNPs cover in a matrix, the F_ROH_ was computed for each animal based on the spread of ROH within the different length classes: 1–2 Mb, 2–4 Mb, 4–8 Mb, 8–16 Mb, as well as >16 Mb. The Identity disequilibrium (ID) was calculated using inbreedR package [18] in R software.

## Results

### Patterns of runs of homozygosity distribution

The study discovered 2226 homozygous segments in the genome of the 87 individuals computed for ROH. The mean ROH per animal was 29.71, with values extending from 2.35 to 85.31. The mean length of the ROH was 5.84 Mb, as well as the longest segment, was 51.43 Mb (1,727 SNPs) found in BTA3. The study found that most ROH lengths are relatively short, and most ROH with lengths ranging from 0.13 to 51.43 Mb, accounting for 68.50% of the total number, while the proportion of the genome covered by them was relatively small (Table 1). The size of ROH varies from 0.13Mb to 51.43Mb, and the study found that the longest ROH was situated on BTA3 (51.43Mb, with 1,727 SNPs), while the shortest ROH was located on BTA15 (0.13Mb, with 56 SNPs).

**Table 1.** Mean length and SNP per ROH and amount of ROH per animal in the Pakistani Sahiwal cattle population.

|  | Mean | Maximum | Minimum |
| --- | --- | --- | --- |
| Length in Mb | 5.84 | 51.43 | 0.13 |
| SNP/Animal | 183.6 | 1727.0 | 50.0 |
| ROH/Animal | 29.7 | 85.31 | 2.35 |

### Runs of homozygosity in the Pakistani Sahiwal cattle population

Fig 1 illustrates the link amid the entire amount of ROH and the entire length of the genome that ROH covers in each individual. This relationship varies substantially from one kind of animal to the next. Within this population, the longest-distance animal with an exceptionally long ROH measured 51.43Mb. There was a large difference between the individuals in the total amount of ROHs and their length. Furthermore, the study observed that the amount of ROH varied between chromosomes, while the highest amount of ROH was detected in BTA6 (157 ROH segments), and the minimum amount of ROH was detected in BTA25 (38 ROH segments) (Fig 2).

**Fig 1.**
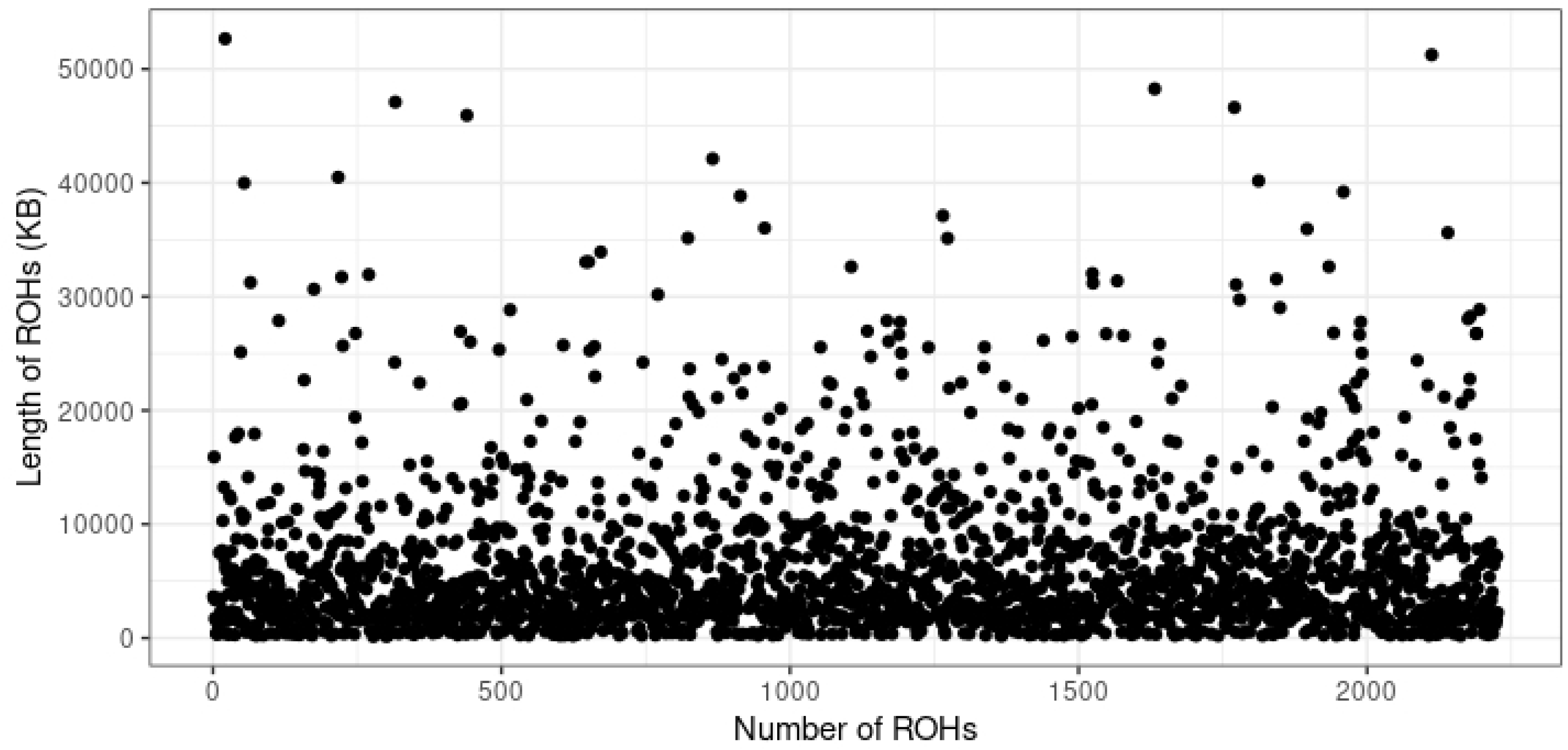
Relationship between the entire number of ROH segments and the entire length (Kb) of the genome in ROH for the entire individuals. Each dot signifies an individual

**Fig 2.**
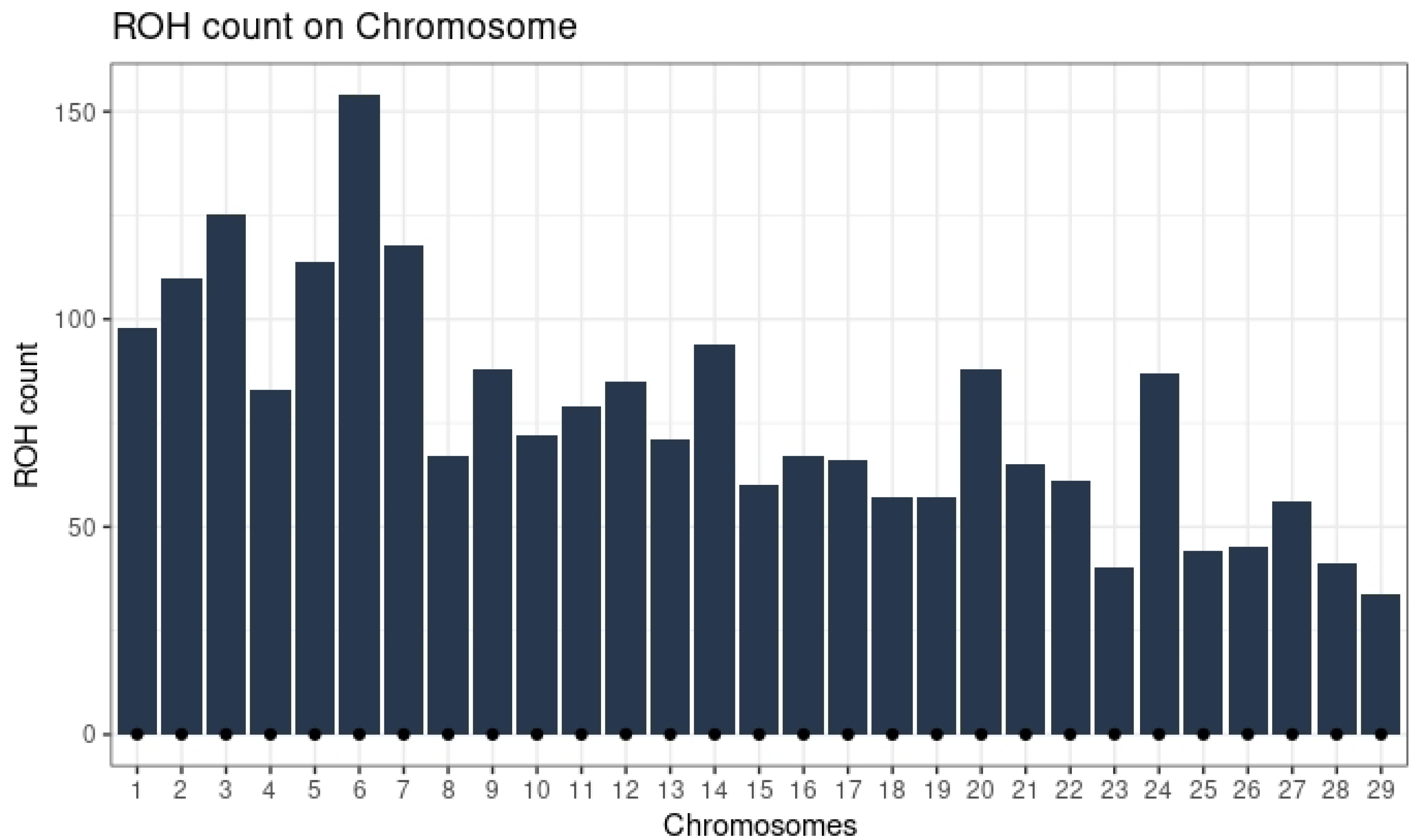
Manhattan plot of the dispersal of runs of homozygosity island in the Sahiwal cattle genome.

### Descriptive statistic of runs of homozygosity amount and length by classes

The different classes of ROH are found on all the chromosomes of the Sahiwal cattle (Fig 3). BTA1 has the highest number of short ROH (1-2Mb), while BTA2 has the highest number of long ROH (>16Mb). The entire length of the ROH for Sahiwal is typically composed of many short and medium segments (2–4 Mb and 4–8 Mb), accounting for approximately 54.18% of all detected ROH (Table 2). On the contrary, the larger ROH greater than 16 Mb was at least a smaller amount, comprising 8.8% of all ROH detected. However, the differences in proportion among the ROH classes are not large, implying that all of the various classes were equally distributed throughout the Sahiwal genome.

**Fig 3.**
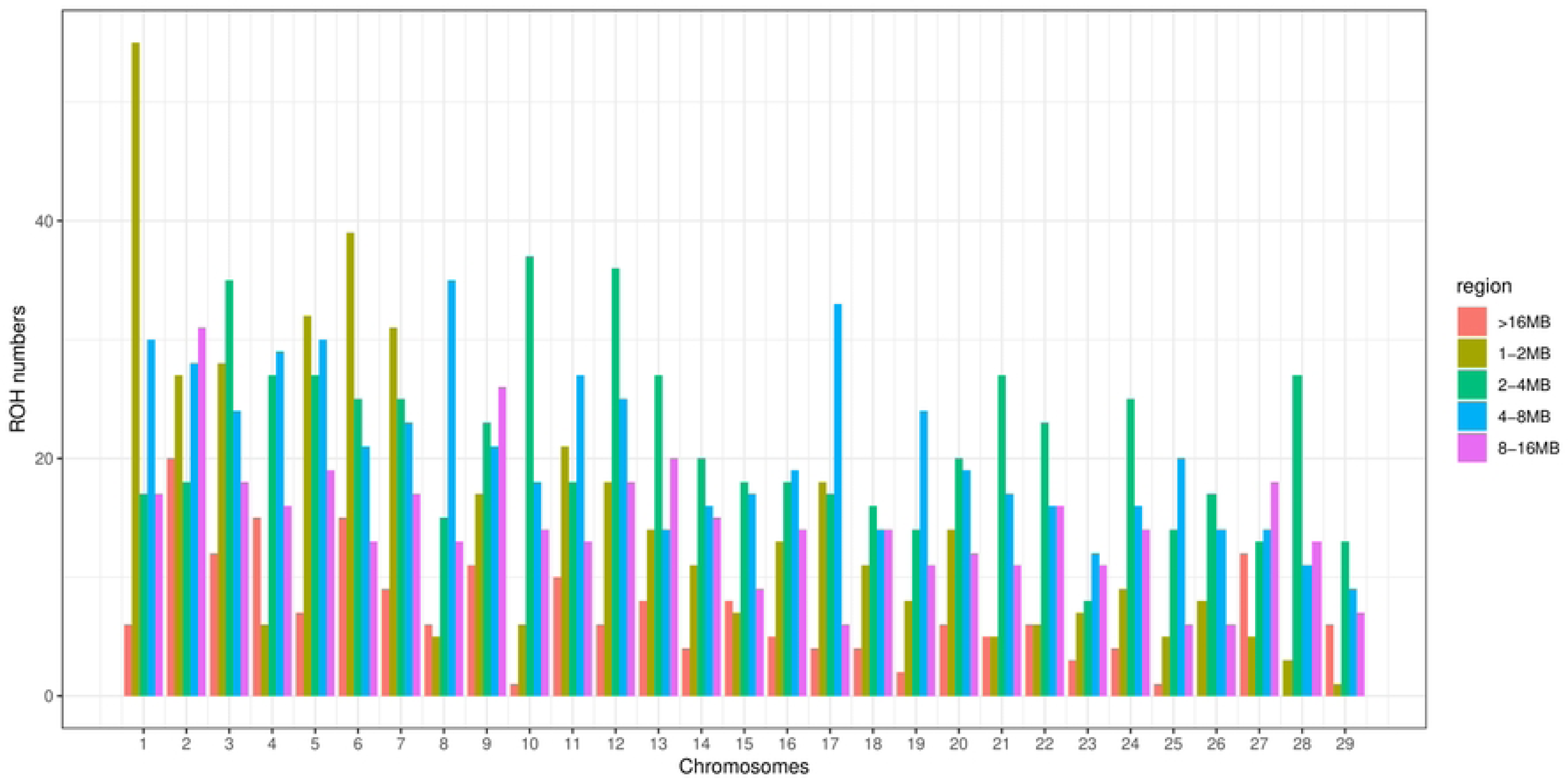
Distribution of different classes of ROH on each chromosome.

**Table 2.**
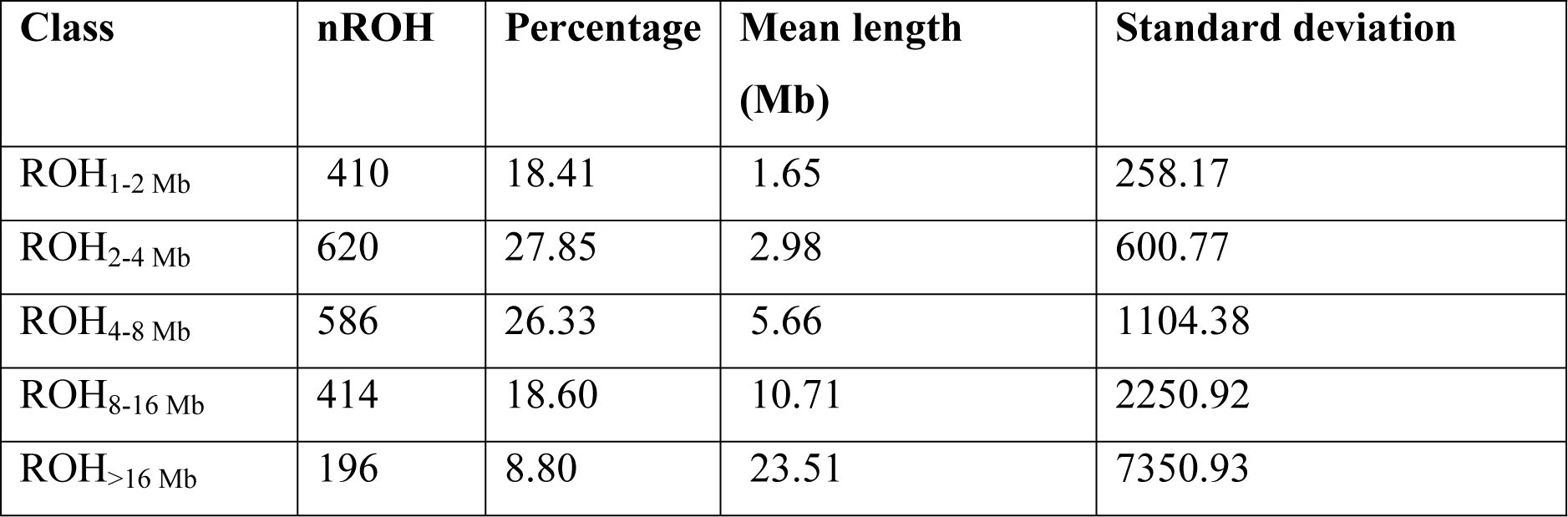
Descriptive statistics of runs of homozygosity amount (nROH) as well as length (in Mb) by ROH length class.

| Class | nROH | Percentage | Mean length (Mb) | Standard deviation |
| --- | --- | --- | --- | --- |
| ROH<sub>1-2 Mb</sub> | 410 | 18.41 | 1.65 | 258.17 |
| ROH<sub>2-4 Mb</sub> | 620 | 27.85 | 2.98 | 600.77 |
| ROH<sub>4-8 Mb</sub> | 586 | 26.33 | 5.66 | 1104.38 |
| ROH<sub>8-16 Mb</sub> | 414 | 18.60 | 10.71 | 2250.92 |
| ROH<sub>>16 Mb</sub> | 196 | 8.80 | 23.51 | 7350.93 |

### Analysis of the inbreeding coefficient (F_ROH_)

Descriptive statistics of inbreeding coefficients based on ROH for different classes of ROH are shown in Table 3. The F_ROH_ coefficient, calculated for all detected ROHs longer than 1 Mb, is one of the most reliable inbreeding coefficients, as it accurately describes the inbreeding phenomena caused by previous and current selection. The mean value of the F_ROH_ coefficient ranged from 0.016 to 0.057. Shorter ROH had the highest mean F_ROH_ values (F_ROH1-2_=0.057), even though they were estimated for ROH greater than 1 Mb. However, the comparatively low F_ROH_ values were typical of medium ROH (F_ROH4-8_=0.016). However, for F_ROH_ computed based on ROH with lengths >16 Mb, the highest values were detected for the coefficient of variation (66.43%).

**Table 3.** Descriptive statistics of the genomic inbreeding coefficients based on runs of homozygosity (FROH) for different Classes of ROH.

| Inbreeding coefficient | Mean | Median | Minimum | Maximum | Coefficient of variation (%) | N |
| --- | --- | --- | --- | --- | --- | --- |
| $F_{ROH1-2}$ Mb | 0.057 | 0.057 | 0.043 | 0.085 | 11.88 | 87 |
| $F_{ROH2-4}$ Mb | 0.017 | 0.017 | 0.007 | 0.033 | 27.58 | 87 |
| $F_{ROH4-8}$ Mb | 0.016 | 0.016 | 0.002 | 0.038 | 47.09 | 85 |
| $F_{ROH8-16}$ Mb | 0.022 | 0.019 | 0.003 | 0.007 | 59.39 | 83 |
| $F_{ROH>16}$ Mb | 0.026 | 0.022 | 0.085 | 0.070 | 66.43 | 69 |

**Table 4.** Characterization of genomic regions with the highest frequency of occurrence of ROH.

| Chromosomes | Start position | End position | Length (MB) | Number of SNPs | Number of genes |
| --- | --- | --- | --- | --- | --- |
| 1 | 81603436 | 85420146 | 3.73 | 50 | 33 |
| 1 | 80349645 | 85371569 | 4.90 | 69 | 46 |
| 1 | 79557477 | 85697413 | 6.00 | 93 | 58 |
| 4 | 62376334 | 68687534 | 6.16 | 94 | 22 |
| 4 | 63005973 | 68090784 | 4.97 | 77 | 18 |
| 4 | 30416555 | 34689529 | 4.17 | 68 | 25 |
| 4 | 10029115 | 13667494 | 3.55 | 60 | 19 |
| 5 | 55015181 | 63604786 | 8.39 | 132 | 80 |
| 5 | 25860120 | 30048658 | 4.09 | 66 | 12 |
| 5 | 56197180 | 62322675 | 5.98 | 99 | 24 |
| 5 | 1.07E+08 | 1.17E+08 | 9.66 | 164 | 31 |
| 8 | 52616884 | 62313669 | 9.47 | 137 | 62 |
| 8 | 52593147 | 58105422 | 5.38 | 88 | 9 |
| 12 | 25702340 | 29019164 | 3.24 | 51 | 10 |
| 12 | 25845412 | 29019164 | 3.10 | 50 | 9 |
| 13 | 47546608 | 51375198 | 3.74 | 60 | 17 |
| 16 | 40838497 | 46587340 | 5.61 | 78 | 71 |
| 18 | 50633635 | 54242897 | 3.52 | 55 | 54 |
| 18 | 57238311 | 60821801 | 3.50 | 58 | 16 |
| 18 | 56824366 | 60821801 | 3.90 | 65 | 19 |
| 18 | 43625758 | 54021201 | 10.15 | 170 | 81 |
| 19 | 41444229 | 48781192 | 7.17 | 86 | 147 |
| 19 | 37870579 | 48933619 | 10.80 | 146 | 163 |
| 19 | 38037897 | 49936700 | 11.62 | 176 | 188 |
| 19 | 44270154 | 51617706 | 7.17 | 121 | 135 |
| 19 | 44216479 | 51680150 | 7.29 | 124 | 108 |
| 21 | 26123921 | 30558507 | 4.33 | 72 | 29 |
| 29 | 41400225 | 45461273 | 3.97 | 66 | 16 |

**Table 5.** Candidate genes associated with milk-related traits in the genomic regions with the highest frequency of occurrence of ROH employing region-based association analysis.

| Chromosome | Start Position | End Position | Length (MB) | Candidate genes |
| --- | --- | --- | --- | --- |
| 2 | 65079608 | 96273076 | 31.19 | **STAT1** |
| 3 | 73035441 | 84336502 | 11.30 | IL12RB2 |
| 3 | 15791927 | 23652002 | 7.86 | ANXA9 |
| 3 | 14943150 | 27642301 | 12.70 | **ATP1A1, ANXA9** |
| 3 | 6418903 | 16216163 | 9.80 | FCGR2B |
| 4 | 56786911 | 96770424 | 39.98 | LEP |
| 4 | 76859 | 27842356 | 27.77 | GRB10 |
| 5 | 99339453 | 1.05E+08 | 5.42 | **OLR1, CD4** |
| 7 | 48812901 | 65930546 | 17.12 | CD14 |
| 8 | 9837754 | 15936813 | 6.10 | CLU |
| 13 | 50339740 | 68706267 | 18.37 | SRC |
| 15 | 30021603 | 63942034 | 33.92 | ZNF215 |
| 15 | 14135810 | 65380895 | 51.25 | BCO2, ZNF215 |
| 17 | 33143310 | 38090810 | 4.95 | IL2 |
| 19 | 37870579 | 48933619 | 11.06 | GH1, STAT5A, STAT5B |
| 19 | 23314971 | 29801929 | 6.49 | ATP1B2 |
| 19 | 48135309 | 53441712 | 5.31 | FASN |
| 19 | 7557250 | 30022677 | 22.47 | CCL2, ATP1B2 |
| 20 | 30116920 | 32986004 | 2.87 | GHR |
| 21 | 13937115 | 49087612 | 35.15 | RASGRF1 |
| 22 | 38727002 | 55892775 | 17.17 | LTF |
| 26 | 15004908 | 23013248 | 8.01 | SCD |

### Effects of inbreeding on heterozygosity and fitness

Fig 3 shows the g2 measure, which serves as a quantitative assessment of identity disequilibrium (ID). This measure can be employed as part of a broader framework to quantify the direct influence of inbreeding on marker heterozygosity and fitness. The study result reveals an average g2 of 0.00114 and a 95% confidence interval of 0.00092 – 0.0013.

Alternatively, the heterozygosity–heterozygosity correlation (HHC) coefficients were computed (Fig 4). The study result reveals an average HHC of 0.957 and a 95% confidence interval of 0.943 – 0.967.

**Fig 4.**
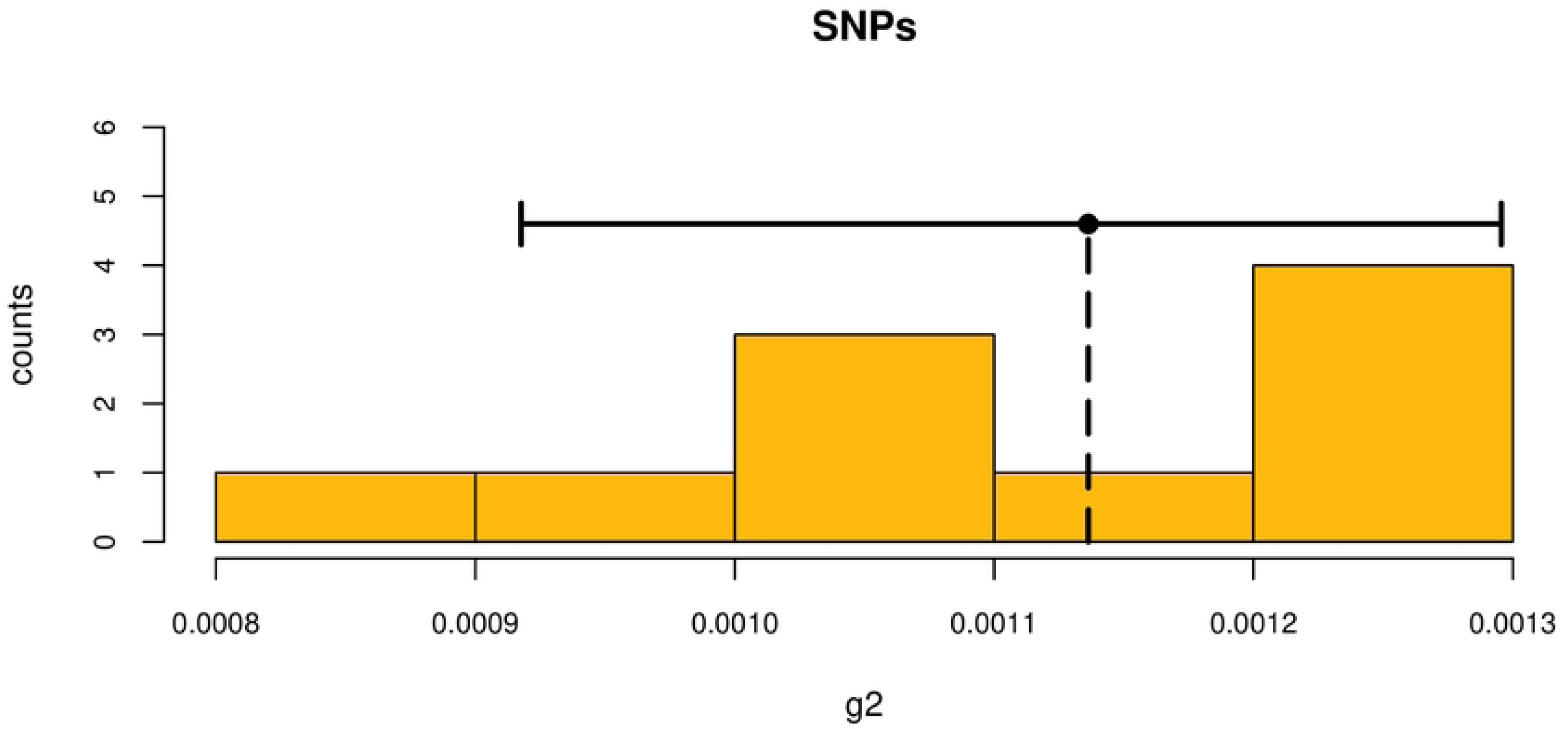
Distribution of g2 from bootstrapping with confidence interval.

**Fig 5.**
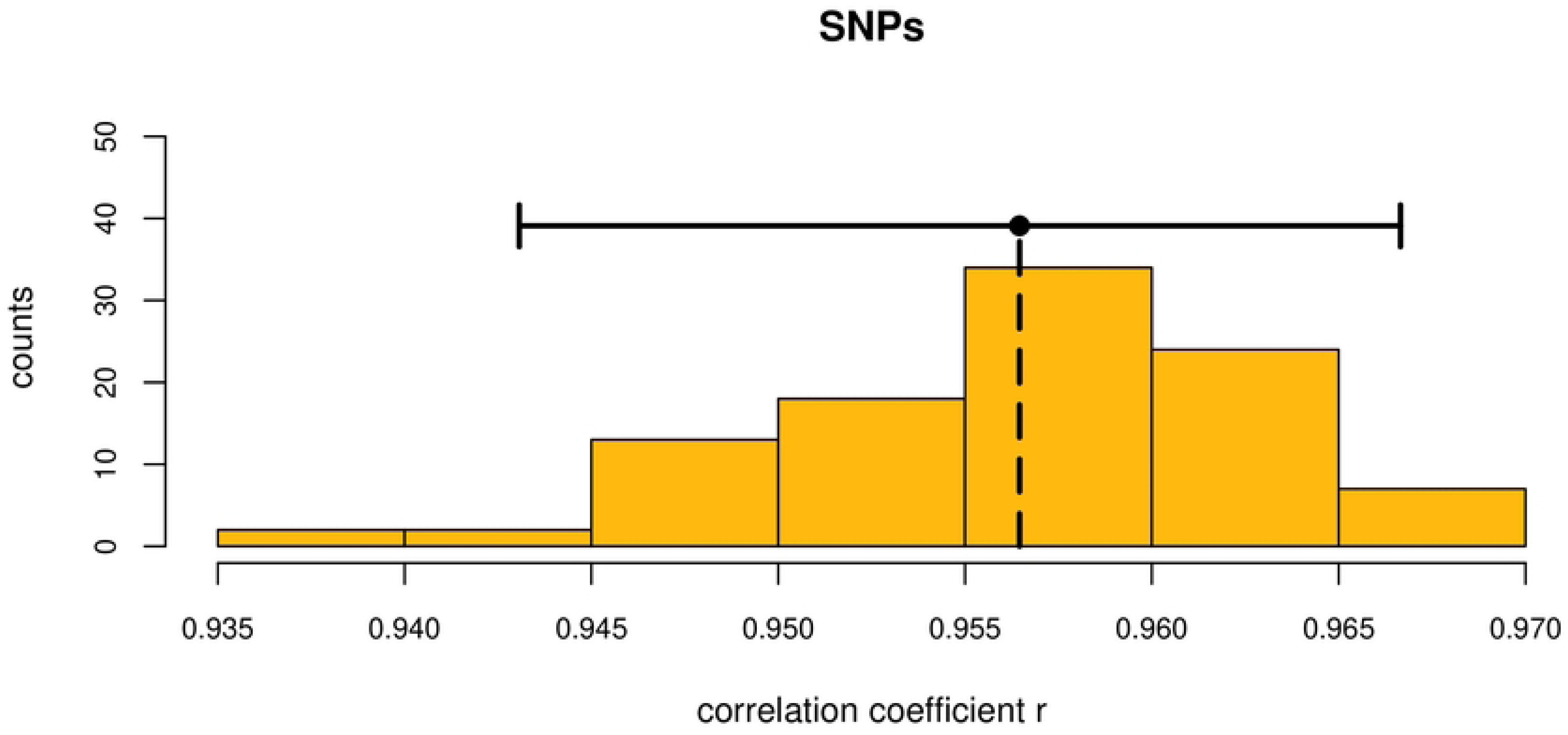
Distribution of the coefficient of heterozygosity-heterozygosity correlations with confidence interval.

### Characteristics of genomic regions with a high frequency of runs of homozygosity occurrence

The ROH was calculated in the 29 autosomes of Sahiwal cattle reared in Pakistan to find genomic areas with the highest incidence of ROH, possibly under the impact of selection. The study then determined the relative occurrence of each SNP in ROH and chose the top 1% of the markers that occur the most frequently. The regions of the genome in which haplotypes are expected to be under selection pressure were indicated by the closest marker with the highest frequency of ROH occurrence. The study identified 28 genome regions with a high incidence of ROH for Sahiwal cattle on BTA 1, 4, 5, 8, 12, 13, 16, 18, 19, 21 and 29. The study observed eight regions with more SNPs on BTA 5, 8,18, and 19.

## Discussion

The study discovered 2,226 ROH in the 87 animals, with a mean of 29.71 ROH per animal. The highest number of ROH per chromosome was described in BTA6. The results are similar to those of Mulim et al. (2022) observed in 16 worldwide cattle populations; BTA1, BTA6, and BTA7 showed the highest ROH concentrations. Similarly, a ROH island was found in BTA6 in Gyr cattle [15]. ROH islands similar to those reported herein Sahiwal were also found on the Brahman, Gyr, and Nellore cattle chromosomes [48]. The ROH islands discovered in BTA6 were also reported in Italian Holstein cattle [49], Tyrol Grey cattle [50], as well as dairy and beef cattle breeds [51]. It is important to note that BTA6 is known to include genes that influence milk yield traits [52], suggesting that signals of selection for dairy qualities may be reflected in high autozygosity in certain chromosomal regions. There was also evidence of overlap between the ROH islands in the cattle breeds originally chosen for beef and dairy production, indicating that the pressure of selection may act on traits unrelated to these two categories. An ROH in BTA6 was shown to be related to fat, protein, and milk production in Mexican Holstein cattle [53]. This ROH was 5.21 Mb in length. [54] found relationships between several QTLs and the milk fat content in Dutch Holstein cattle. The US Holstein cattle were studied by [55], and they found several relationships and QTLs for conformation and productive qualities in the same genomic region.

In the study, BTA3 contained the longest ROH among Pakistani Sahiwal cattle at 51.43 Mb. Similar results have been obtained in Jersey dairy cattle; [56] identified two regions with higher ROH frequencies in all populations on BTA3 and BTA7, between 42.4 and 44.2 Mb and 35.6 and 41.4 Mb, respectively. [57] determined the longest ROH for BTA3 in Chinese Hainan cattle. [58] and [59] reported much higher results on BTA8 in a contemporary Holstein heifer (87.13 Mb) and a Cinisara cattle breed (112.65 Mb). Contrary to the study findings, [60] found Chinese Wagyu Beef Cattle with the longest ROH on BTA10 (48.6 Mb) as well as the shortest ROH in BTA3 (0.5 Mb).

Most of the ROH length in the Pakistani Sahiwal cattle population comprised short and medium segments (2–4 Mb and 4–8 Mb, respectively). These segments represented approximately 53.88% of all ROHs identified in the study. [61] also discovered that the entire length of ROH in the Shanghai Holstein cattle population consisted primarily of shorter segments (1–2 Mb and 2–4 Mb). Such segments reported for almost 78% of the entire ROH discovered and donated between 0.79 and 2.16 percent to the entire length of the ROH. [62] discovered that medium ROH 4 to 8 Mb, as well as long ROH greater than 8Mb segments, predominated in the genome of Montana beef cattle, accounting for approximately 74% of all ROH discovered and contributing significantly to 90% of the collective length of ROH. They stated that the high amount of medium and long ROH might result from the inability of low-density arrays to recognize ROH between 0.5 and 2 Mb in length.

However, these results contradict some of the earlier reported findings in cattle [6,15,63], sheep [40], as well as pigs [64], in which the entire length of ROH was typically composed of a large amount of shorter ROH. It is important to emphasize that comparing ROH studies is difficult because the parameters used to define ROH do not consistently align. The absence of agreement makes it possible for various thresholds to be used in different research [65], and it may cause bias in ROH-based autozygosity estimates [6]. Most of the above research used medium-density arrays (50K), indicating much shorter ROH. According to [6], the 50K array tends to expose many short segments. Inbreeding that occurred in the distant past is reflected in ROH segments that are short, whereas ROH segments that are lengthy are often generated by inbreeding that occurred in more recent times [50,66,67]. According to [68], the occurrence of segments bigger than 10 Mb may be traced back to inbreeding from currently shared descendants just five generations ago. This revealed that inbreeding events from both the distant past and more recent times could have affected this population, although the genome of the Sahiwal cattle population has been mostly altered by inbreeding or selection pressures that have occurred more recently [41,69].

In the study, the inbreeding coefficient based on ROH (F_ROH_) ranged from 0.016 to 0.057. Various studies have documented a diverse range of estimates for the level of inbreeding coefficient based on ROH (F_ROH_) in different domestic animal species. In a recent study conducted by [70], an investigation was carried out to assess the level of inbreeding based on runs of homozygosity (F_ROH_) in different groups of Pakistani cattle breeds. The results revealed varying levels of inbreeding within each group. Specifically, the dairy group exhibited a range of F_ROH_ values of 0.010 to 0.078, the Dual group showed a range from 0.002 to 0.057, and the Draft group displayed a range from 0.006 to 0.078. [6] conducted a study in which they observed a range of 0.087– 0.156 for F_ROH_ > 1 Mb among four different cattle breeds. In their study, [42] documented a spectrum of values ranging from 0.042 to 0.113 for F_ROH_ >1 Mb among eight distinct cattle breeds. A comparable estimated range of 0.026–0.190 for F_ROH_ >1 Mb was documented in six different breeds of goat in China [71]. [72] reported the observation of a range of 0.002 – 0.344 in the Chinese composite pig breed for F_ROH_ > 1 Mb. [67] reported an estimated range of 0.016 – 0.099 for F_ROH_ >1 Mb in different breeds of sheep in Italy. Unequal marker densities employed in different research may contribute to variations in the estimations of inbreeding rates derived from ROH [73].

The study observed an average HFC of 0.00114 which represents the correlation between genetic heterozygosity and fitness, indicating the impact of inbreeding on health and viability. In a scientific context, a positive HFC value within this confidence interval means that maintaining genetic diversity (heterozygosity) is essential to maintaining the health and productivity of the Sahiwal cattle population [74,75]. Higher heterozygosity contributes to improved fitness, reducing the risk of inbreeding depression and promoting long-term viability [76]. The correlations between heterozygosity and fitness have been studied extensively in animal populations due to advances in molecular techniques [74,77]. Although several investigations have discovered limited or inconclusive associations between heterozygosity and fitness, alternative research has shown notable connections, underscoring the need to understand the fundamental mechanisms and genetic processes that influence HFCs in many populations [74,78,79].

The study revealed an average heterozygosity–heterozygosity correlation coefficient (HHC) of 0.957, which indicates a strong positive correlation between genetic heterozygosity and genetic diversity within the Sahiwal cattle population [23,80]. The positive HHC coefficient underscores the importance of preserving genetic diversity to prevent inbreeding and its associated negative effects on population health and productivity [81]. Maintaining genetic diversity is crucial for the long-term health and adaptability of the Sahiwal cattle population [82–84].

The study revealed several genes that are associated with milk production traits in the Sahiwal cattle. The Signal Transducer and Activator of Transcription 1 (STAT1) detected on BTA2 in the study has been reported in previous studies to be associated with milk production traits [85,86]. [87] reported the involvement of STAT1 in regulating gene transcription related to milk protein synthesis and fat metabolism. The STAT1 gene has been identified as one of several genes empirically shown to impact the expression of production traits in cattle [88]. The study conducted by [89] found a significant association between the STAT1 gene and female fertility in Nellore cattle, along with several other genes. The STAT1 gene has been identified as a potential candidate associated with the age at first birth in cattle, primarily due to its functional properties [90]. According to [91], the interplay of SNPs within the STAT3 gene and the interactions between the STAT1 and STAT3 genes have a substantial role in the observed diversity of embryonic survival in cattle.

The Sodium/potassium-transporting ATPase subunit alpha-1 (ATP1A1) detected on BTA3 in the study has been found to be associated with milk production traits. The study by [92] revealed that ATP1A1 significantly influenced the milk protein percentage in Holstein dairy cows. The study conducted by [93] found a significant association between ATP1A1 and both Heart Girth (HG) and Hip Height (HH) in Chinese Holstein cattle. Several studies have identified ATP1A1 as a potential candidate gene associated with thermotolerance in dairy cattle, making it a potential marker for selecting individuals with enhanced heat tolerance [94–97].

The oxidized low-density lipoprotein receptor 1 (OLR1) detected on BTA5 in the study has been found to be associated with milk production traits. The study by [98] showed a noteworthy correlation between the OLR1 SNP and protein percentage and somatic cell score. Furthermore, a study by [99] revealed a statistically significant correlation between OLR1 gene expression and milk fat percentage in Dutch Holstein-Friesian cattle. According to [100], the OLR1 gene’s genetic polymorphism notably influenced the milk fat content in Holstein Friesian dairy cows. According to [101], the OLR1 area exhibited a statistically significant impact on the milk fat percentage and milk fat output. Based on the fact that OLR1 serves as a receptor for oxLDL and shows high expression levels in cardiac tissues, it is plausible that OLR1 could exert a direct influence on the metabolism of oxLDL, thus impacting lipid metabolism [101]. According to [102], a substantial association was seen between the OLR1 marker and various phenotypic traits in Simmental cattle, including final weight (FW), hot carcass weight (HCW), chilled carcass weight (CCW), dressing percentage (DP), and total weight gain (TWG).

The cluster of differentiation 4 (CD4) detected on BTA5 in the study has been found to be associated with milk production traits. The study by [103] revealed a strong association between SNPs in the bovine CD4 gene and fat percentage. [104] reported a strong association between SNPs in the bovine CD4 gene and the annual milk output and the prevalence of mastitis in dairy cattle. In their study, [105] found a strong association between SNPs in the bovine CD4 gene and the estimated breeding values (EBVs) for milk yield, protein yield, and somatic cell score (SCS) in Chinese Holstein cows. According to the findings of [106], there was a substantial association between SNPs in the bovine CD4 gene and both the frequency of clinical mastitis occurrence and annual milk output. The findings of their study indicate that the CD4 gene should be regarded as a potential candidate gene, and the found SNPs may serve as valuable molecular markers to select dairy animals that possess genetic resistance to the development of mastitis.

## Conclusion

Sahiwal cattle are Pakistan’s most important breed of dairy cattle due to their tolerance to heat, resistance to disease, and adequate performance in low-quality roughages. The ROHs were identified in all animals. The entire length of the ROH for Sahiwal is typically composed of many short and medium segments (2–4 Mb and 4–8 Mb), accounting for approximately 54.18% of all detected ROH. The inbreeding coefficient based on ROH (F_ROH_) ranged from 0.016 to 0.057. The study observed an average HFC of 0.00114 and an HHC of 0.957. The study revealed several genes, including STAT1, ATP1A1, OLR1, and CD4, some known essential genes related to milk production. Thus, understanding ROH patterns, an inbreeding coefficient derived from ROH, and candidate genes associated with significant economic traits can help explain the mechanisms governing these traits in Sahiwal cattle. Furthermore, the results obtained could provide the groundwork for future research on economically significant traits associated with milk production in Sahiwal cattle.

## Author contributions

A.R.S. and M.S.R. designed the study, analyzed the data and wrote the drafted manuscript. F.R. and F.S.A. interpreted the data. All the authors reviewed the paper. All authors have read and agreed to the current version of the manuscript.

